# *Muilla lordsburgana* (Asparagaceae: Brodiaeoideae), a new species found north of Lordsburg, southwestern New Mexico

**DOI:** 10.1101/2020.09.24.312215

**Authors:** Patrick J. Alexander

## Abstract

*Muilla lordsburgana* P.J.Alexander sp. nov. is described from eastern Lordsburg Mesa in the northwestern fringe of the Chihuahuan Desert, southwestern New Mexico. It is very similar to *Muilla coronata*, a species known in the Mojave Desert of California and a small area of adjacent Nevada. Compared to *Muilla coronata*, the style and stigma combined are longer, the anthers are longer, the fruits are larger, and the seeds are larger. It has a short flowering period in March, and is difficult or impossible to find at other times of the year. It is narrowly distributed, limited to a band of deep, coarse, sandy soils derived primarily from granite. Several species that are otherwise uncommon in the area also occur in this habitat, including *Logfia depressa, Pectocarya platycarpa*, and *Plagiobothrys arizonicus*. Invasive species, especially *Erodium cicutarium* in the winter annual flora, are generally dominant. *Muilla lordsburgana* has probably been overlooked until now because it is not easy to observe and occurs in habitats that are not attractive to botanists.

---

On 15 Mar 2015, I happened upon a small geophytic monocot north of Lordsburg that was unknown to me. I identified it as *Androstephium breviflorum* S.Watson. However, in February 2017 after encountering genuine *Androstephium breviflorum* in Arizona, I realized this was an error. Searches in increasingly distant regional floras led to *Muilla coronata*. It was immediately obvious that the plant north of Lordsburg was very closely related to this species, and possibly conspecific. However, *Muilla coronata* is known only in the Mojave Desert in California and ±60 km into Nevada near Las Vegas. Its closest populations are ±700 km distant from Lordsburg. Although such disjunctions do occur, they are unusual. So, in March 2017 I set out to further document these plants and determine if they are conspecific with *Muilla coronata*.

*Muilla coronata* is a very distinctive species. There are few geophytic monocots with petaloid filaments in the southwestern United States. Within this small group, *Muilla coronata* is the only one with 6 equal, free stamens and no obvious hypanthium. The others have stamens with the filaments connate (*Androstephium* Torr.), stamens obviously unequal or some reduced to staminodes (*Brodiaea* Sm., *Dichelostemma* Kunth, *Dipterostemon* Rydb.), an obvious and well-developed hypanthium (*Triteleia* Douglas ex Lindl.), or several of these features. Perhaps because it is so distinctive, descriptions of *Muilla coronata* have often been cursory (Greene 1888; Jepson 1909; Abrams & Ferris 1923; Ingram 1953; Munz & Keck 1959). More recent descriptions have provided more detail (Shevock 1984; Pires & Reveal 2002; Pires 2017). For morphological details of *Muilla coronata* not addressed in these descriptions, I relied on assistance from botanists familiar with *Muilla coronata* and online photographs of *Muilla coronata* (primarily through CalPhotos and iNaturalist). Because the Lordsburg *Muilla* appears to be a rare plant, I collected relatively few specimens and rely heavily on photographs (taken at a 1:1 magnification ratio) for floral measurements.

By late March, after a few days chasing the Lordsburg *Muilla* in the field and working with photographs, two floral distinctions between it and *Muilla coronata* became apparent. The combined style + stigma of my plants (≥2.5 mm) appeared to be longer than that of *Muilla coronata*, based on online photographs. Andrew Sanders confirmed, from specimens at UCR, that the style + stigma are 1–2 mm in *Muilla coronata*. The anthers are also longer, 1.3–1.6 mm in the Lordsburg *Muilla*, 0.6–0.9 mm in *Muilla coronata* (based on Pires & Reveal 2002). When mature fruits and seeds were available, these also turned out to be larger than those of *Muilla coronata*. Four non-overlapping characters seem sufficient basis for a species, particularly given the geographic disjunction between it and *Muilla coronata*, so I here name it *Muilla lordsburgana*.

## Muilla lordsburgana P.J.Alexander, sp. Nov

> (Figs. 1–4) Type: New Mexico. Hidalgo County: Eastern Lordsburg Mesa, 7.2 miles northeast of Ninemile Hill, 4.5 miles southeast of Lone Mountain, and 9.9 miles northwest of Apache Mountain, 32.53363°N 108.70569°W (this & all subsequent coordinates in nad83), elev. 4670 ft., 19 Mar 2017, *Alexander 1566* (holotype: NMC; isotypes: ASU, UCR).

**Figure 1.**
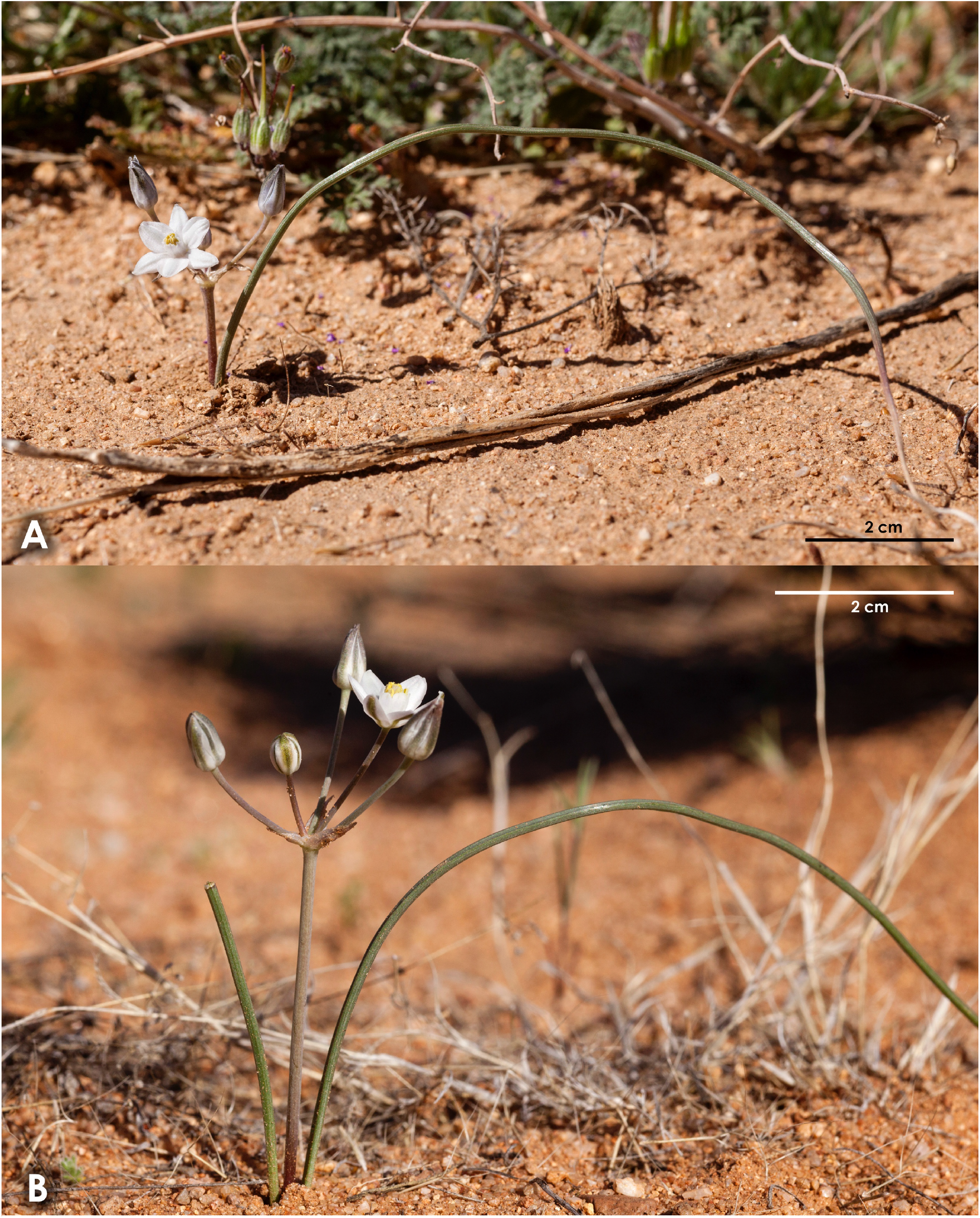
*Muilla lordsburgana* at: **A**, 32.4686°N 108.6456°W, 17 Mar 2017, site of *Alexander 1565*; **B**, 32.5336°N 108.7057°W, 19 Mar 2017, site of *Alexander 1566*.

**Figure 2.**
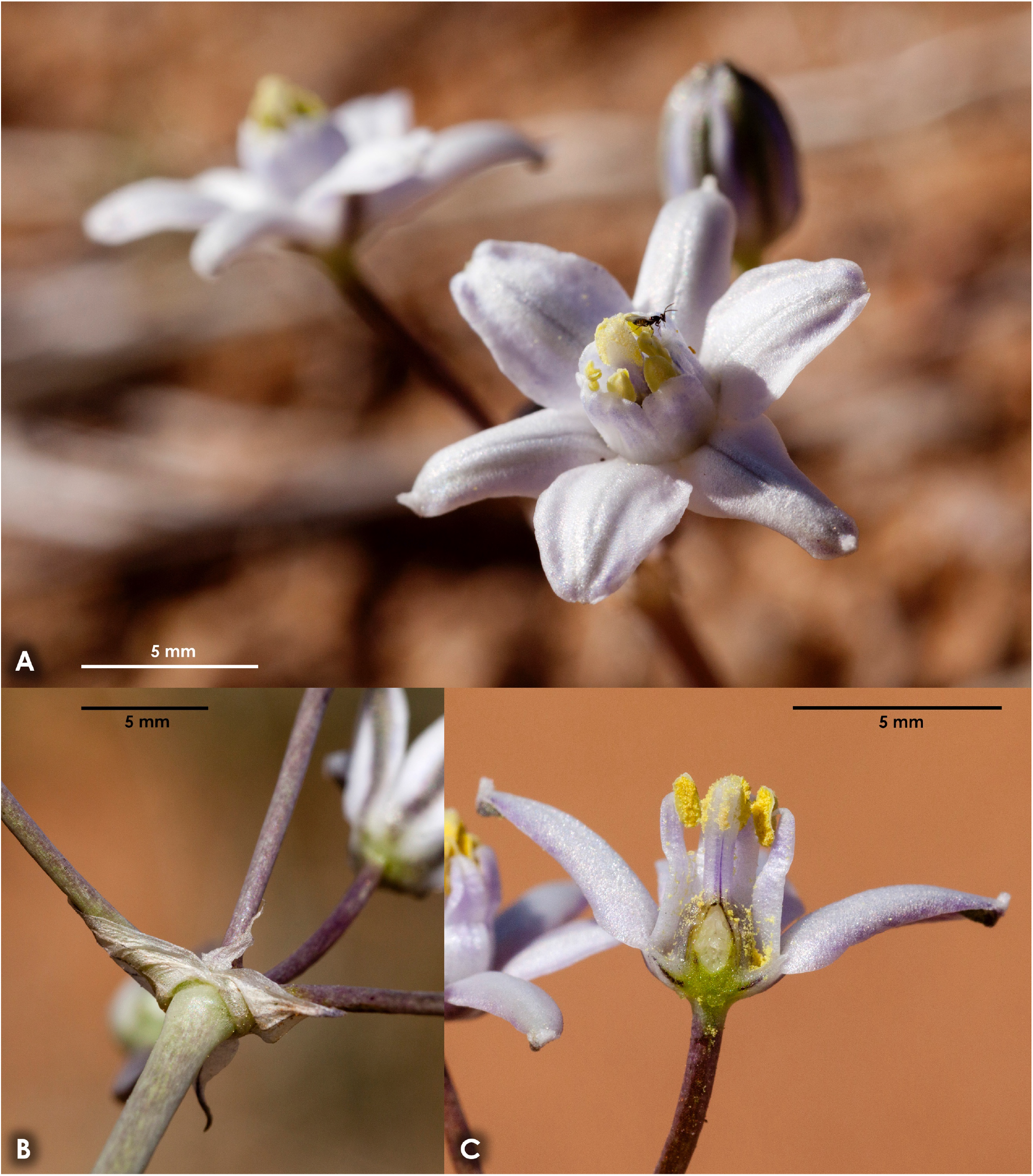
**A**, flowers of *Muilla lordsburgana* at 32.5923°N 108.7302°W, 10 Mar 2017, site of *Alexander 1564*; **B**, bracts at 32.4684°N 108.7055°W, 19 Mar 2017; **C**, longitudinal section of a flower at 32.4823°N 108.7309°W, 18 Mar 2017.

**Figure 3.**
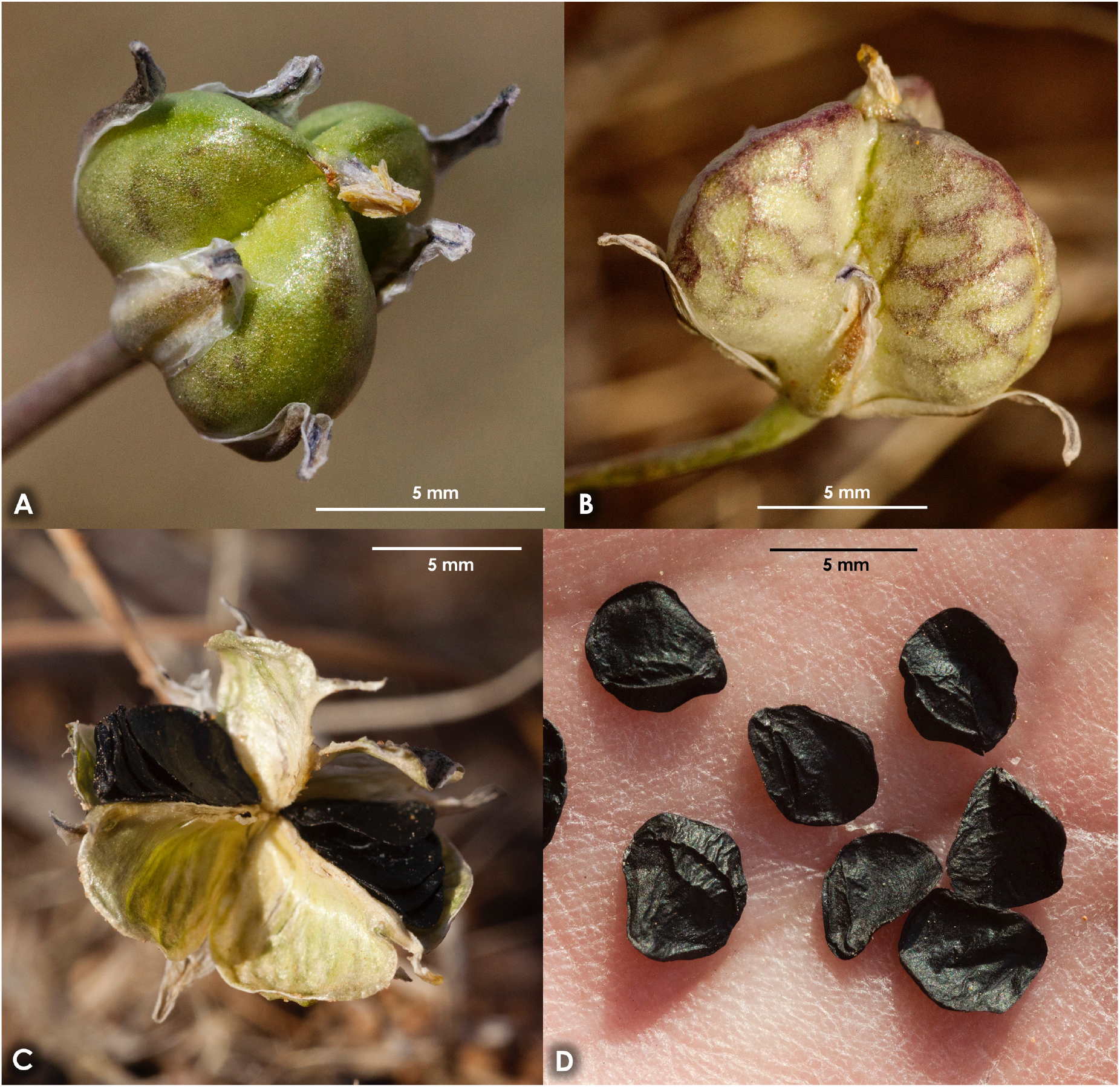
Fruits and seeds of *Muilla lordsburgana*: **A**, immature fruit at 32.4684°N 108.7055°W, 19 Mar 2017; **B**, mature fruit at 32.4686°N 108.6456°W, 28 Mar 2017, collection site of *Alexander 1565*; **C**, dehisced fruit at 32.5133°N 108.7016°W, 7 Apr 2017; D, seeds at 32.5133°N 108.7016°W, 7 Apr 2017.

**Figure 4.**
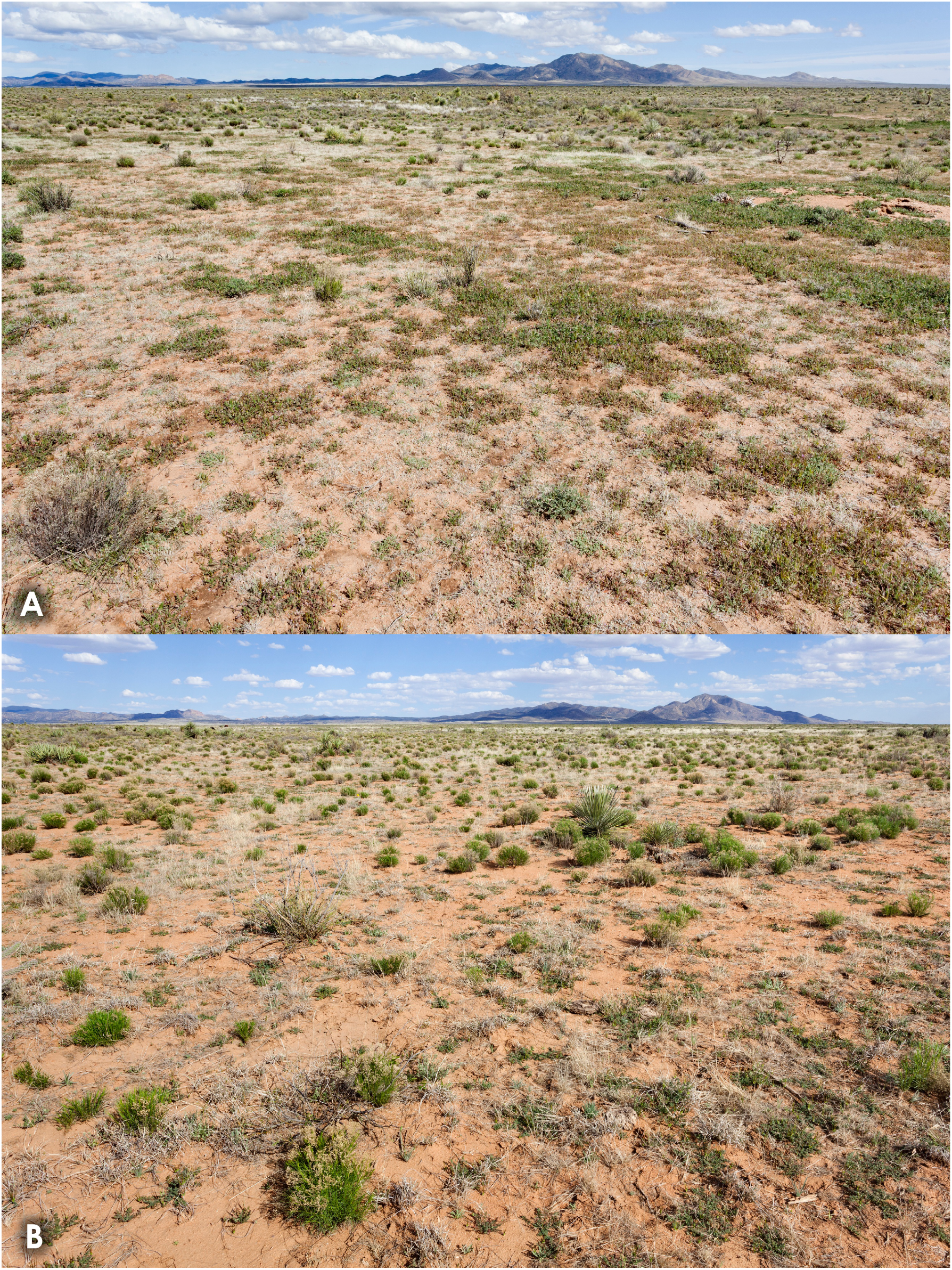
Habitat of *Muilla lordsburgana*: **A**, at 32.47096°N 108.68991°W, 19 Mar 2020; **B**, at 32.51652°N 108.67966°W, 18 Mar 2017.

Similar to *Muilla coronata* Greene but differing as follows: style & stigma longer (combined length ≥2.5 vs. ≤2 mm); anthers longer (≥1.3 vs. ≤0.9 mm); fruits larger (>8 vs. ≤7 mm); seeds larger (>4 vs. ≤3 mm); occurs in the northwestern fringes of the Chihuahuan Desert, ±700 km east-southeast of *Muilla coronata*.

Perennial, scapose, glabrous herbs with fibrous-coated corms. Corms (3.5–)5.5–7(–9) cm below ground, 1.1–1.7 cm wide. Leaves 1–2, basal, withering soon after anthesis, surrounded by a fibrous sheath for 2–4.5 cm above the corm; blade linear, above-ground portion 10–19 cm long, 1.5–2 mm wide, drying to ±1 mm wide, ± succulent, weakly channeled. Scape 1 or rarely 2, cylindrical, above-ground portion (1–)2.5–7(–9) cm high, ±1.5 mm wide, drying to ±1 mm wide. Inflorescences umbellate, terminal, 2–6-flowered; subtended by 3–6 hyaline, lanceolate to triangular bracts in two series, the outer 4.1–8.8 mm × 2.2–5.2 mm, 3–5-nerved, the inner 1.5–4.8 mm × 0.4–1.5 mm, obscurely 2-or 3-nerved; pedicels 0.5–2 cm at anthesis, lengthening in fruit. Perianth subrotate; tepals 6, oblong to broadly lanceolate, connate 0.2–0.5 mm at the base, 5.5–7 mm long, the outer 2.1–3.4 mm wide, the inner 3.1–5.5 mm wide, white to pale lavender adaxially, abaxially white to pale lavender with a prominent dark green midvein, margins of the midvein often purple, the purple color rarely extending outward half or more of the distance to the margins of the tepals. Androecium of 6 erect, epipetalous stamens inserted 0.2–0.5 mm above the base of the tepals; filaments 3.3–5 mm long and 1.4–2 mm wide, petaloid, white to pale lavender, dilated their entire lengths, distinct but their margins appressed to slightly overlapping and forming a tube, apex rounded, truncate, or retuse, with a narrow introrse acuminate extension holding the anther; anthers versatile, yellow, 1.3–1.6 mm long, appressed to the stigma and usually adhering to it after anthesis. Gynoecium syncarpous, 3-carpellate; ovary superior, sessile, broadly ovate, 1.6–2.2 mm long; style and stigma 2.5–3.8 mm long, poorly differentiated; style cylindrical, 1.6–2.3 mm long; stigma truncate-conic to clavate, rounded at the apex, 0.6–1.5 mm long, with three poorly defined stigmatic lines extending downward and becoming impercetible as the stigma joins the style. Fruit capsular, loculicidal, globose and 3-lobed, 8.5–10.5 mm long, pale green, turning tan at dehiscence, with a purplish or dark green line along the center of each carpel and purplish or dark green veins curving inward. Seeds black, obovate to orbicular, flattened, 4.5–6.5 mm long and 3.8–5.2 mm wide, surfaces irregularly angled.

### Paratypes

U.S.A. New Mexico. Hidalgo County: eastern Lordsburg Mesa, on the east side of NM Hwy. 464, 2.6 miles northeast of Lone Mountain between Willow Draw and South Fork Corral Canyon, 32.59227°N 108.73019°W, elev. 4520 ft., 10 Mar 2017, *Alexander 1564* (NMC, UCR); eastern edge of Lordsburg Mesa in Wood Canyon, 4.8 miles west-northwest of Apache Mountain and 8.1 miles east of Ninemile Hill, 32.46858°N 108.64561°W, elev. 4760 ft., 17 Mar 2017, *Alexander 1565* (NMC, UNM).

### Etymology

The specific epithet refers to the town of Lordsburg, which is not often honored. The generic epithet reverses *Allium*, a genus to which this species bears little resemblance.

### Common Name

”Muilla” is often used for *Muilla*, but “noino” is the English equivalent of *Muilla*. I suggest “Lordsburg noino”.

### Phenology and pollination

Flowering in March, from about the 5th to the 31st, fruiting in April. The flowers are visited by flies in the family Bombylliidae, butterflies in Lycaenidae subfamily Polyommatinae, and chalcid wasps, probably in Eulophidae subfamily Tetrastichinae.

### Abiotic ecology

*Muilla lordsburgana* occurs at 4430–4960 ft. elevation on the gentle western and southwestern slopes where the bajada of the Big Burro Mountains meets the flatter topography of Lordsburg Mesa (Fig. 5). The soils are derived primarily from Burro Mountain granite (Gillerman 1964) and consist mostly of two soil map units, Mohave sandy clay loam, 0–5 percent slopes and Sonoita-Yturbide complex (Cox 1973). These are very deep soils, without a root-restrictive layer within 60 in. of the surface. The upper layers, to a depth of ±11 inches, are sandy loams, gravelly sandy loams, or loamy sands. Lower layers are gravelly sandy loams, gravelly loamy sands, gravelly loamy coarse sands, gravelly coarse sands, sandy loams, clay loams, or loams. Based on what I can see of surface soils and infer from soil maps, occupied habitat is limited to the west by aeolian fine sand sheets overlying clay loams (Berino loamy sand, hummocky and Pintura-Berino complex, eroded, in Cox 1973) and to the east by gravelly or cobbly (at least in the upper horizon) loam or clay loam higher on the bajada of the Big Burro Mountains (generally corresponding with the Forrest loam of Cox 1973). To the north it ends abruptly at the breaks dropping to the Gila River. Similar habitat extends southeast without an apparent change in topography or soils, but *Muilla lordsburgana* extends only slightly beyond NM Hwy. 90. The band of occupied habitat is ±18 km north-northwest to south-southeast and ±8 km east to west. Within this area, it is most abundant near shallowly incised, ephemeral drainages.

**Figure 5.**
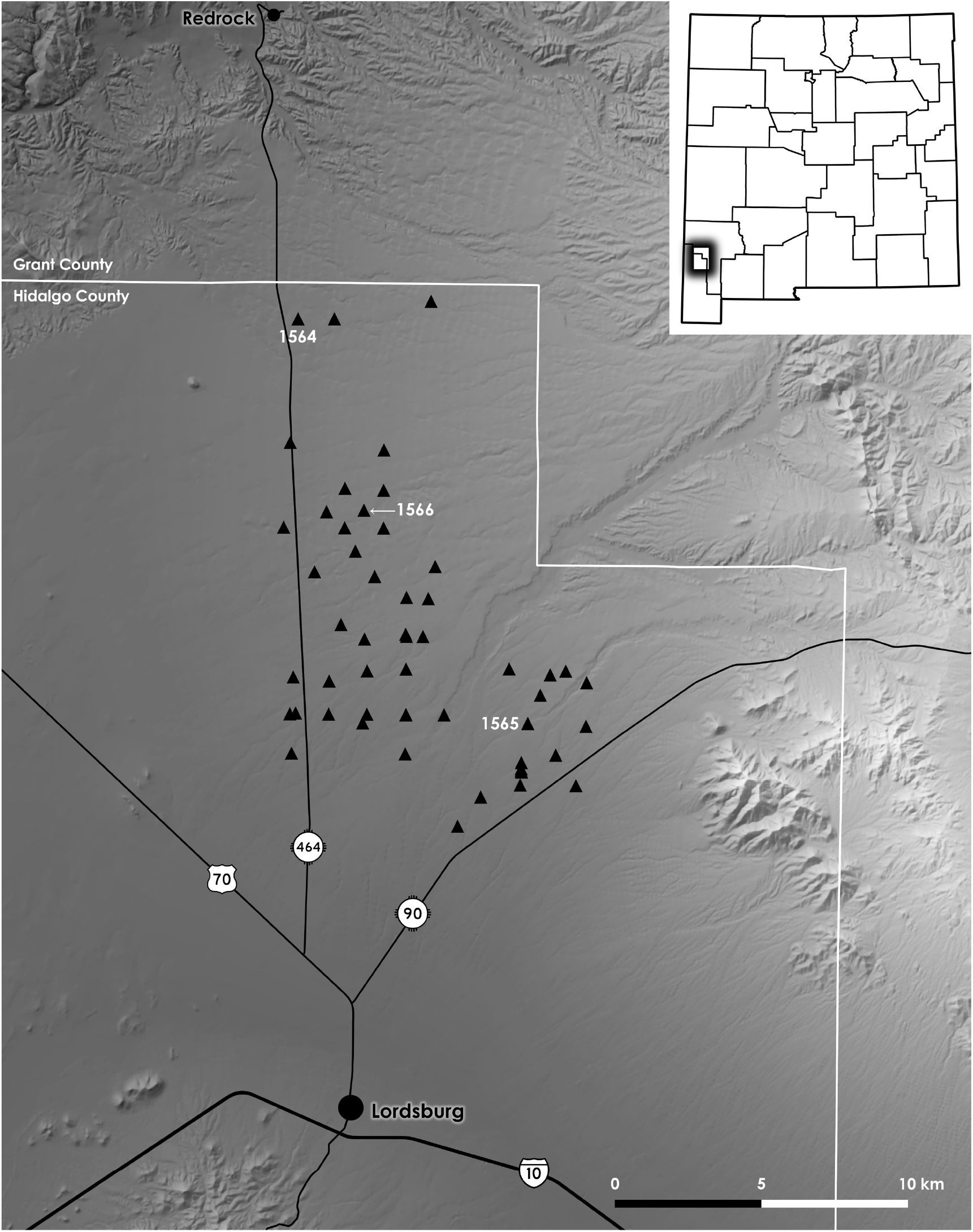
Distribution of *Muilla lordsburgana* (black triangles), with sites of *Alexander 1564, 1565*, and *1566* marked. This and the next map were generated in QGIS (QGIS Development Team 2020).

### Associated plants

The habitat occupied by *Muilla lordsburgana* is generally dominated by annuals, principally by *Erodium cicutarium* (L.) L’Hér. ex Aiton among the winter annuals, *Aristida adscensionis* L. and *Bouteloua aristidoides* (Kunth) Griseb. among the monsoon annuals. In some areas it is dominated by subshrubs, usually *Gutierrezia sarothrae* (DC.) A.Gray, sometimes *Isocoma tenuisecta* Greene. Occasionally it is dominated by perennial grasses, either *Bouteloua eriopoda* (Torr.) Torr., *Eragrostis lehmanniana* Nees, or *Aristida ternipes gentilis* (Henrard) Allred, or by shrubs, *Prosopis glandulosa* Torr. or *Ephedra trifurca* Torr.

Several species occurring with *Muilla lordsburgana* are uncommon in southwestern New Mexico. These include *Plagiobothrys arizonicus* (A.Gray) Greene ex A.Gray, *Logfia depressa* (A.Gray) Holub, *Pectocarya platycarpa* (Munz & I.M.Johnst.) Munz & I.M.Johnst., *Linanthus bigelovii* (A.Gray) Greene, *Leptosiphon chrysanthus* J.M.Porter & R.Patterson, *Nuttallanthus texanus* (Scheele) D.A.Sutton, and *Xanthisma gracile* (Nutt.) D.R.Morgan & R.L.Hartm. These, and others not listed, are significantly associated with *Muilla lordsburgana* and can be used as predictors of suitable habitat beyond the brief window in which it is detectable. This topic will be explored further in another paper, to be published later.

### Habitat condition and potential threats

The historical reference state for the habitat occupied by *Muilla lordsburgana* was probably grassland dominated by *Bouteloua eriopoda*. Cox (1973) identifies these soils as the Sandy and Deep Sand range sites, equivalent to current Sandy (R042XB012NM) and Deep Sand (R042XB011NM) ecological sites (National Resources Conservation Service et al. 2020). These ecological sites are based more on aeolian than alluvial sandy soils, but are nonetheless the best available information on the ecological dynamics of these soils. The historical reference state is perennial grassland for both ecological sites. On the Sandy ecological site, this grassland was dominated by *Bouteloua eriopoda*, with *Sporobolus contractus* Hitchc., *Sporobolus cryptandrus* (Torr.) A.Gray, & *Sporobolus flexuosus* (Thurb. ex Vasey) Rydb. often subdominant, *Muhlenbergia porteri* Scribn. ex Beal & perennial *Aristida* L. also common. Grasslands on the Deep Sand ecological site were historically dominated by *Sporobolus contractus, Sporobolus cryptandrus, Sporobolus flexuosus, & Sporobolus giganteus* Nash, with *Bouteloua eriopoda* or *Muhlenbergia porteri* sometimes subdominant. On both ecological sites, grazing can cause a loss of perennial grasses. On aeolian sands, this is followed by a transition to *Prosopis glandulosa* shrubland, usually with well-developed coppice dunes. Abundant *Gutierrezia sarothrae* is also mentioned as an indicator of grazing-related degradation on the Sandy ecological site. Given that the habitat occupied by *Muilla lordsburgana* has little perennial grass (*Bouteloua eriopoda* is often present, but rarely dominant) while *Gutierrezia sarothrae* is ubiquitous and often abundant, presumably it has been heavily modified from its pre-grazing state. However, while *Prosopis glandulosa* coppice dunes are the typical result on aeolian sands, *Prosopis glandulosa* is sparse on these alluvial granitic sands and coppice dunes at most very poorly developed.

Invasive species are ubiquitous, particularly *Erodium cicutarium & Eragrostis lehmanniana*, as mentioned above, and *Schismus barbatus* (Loefl. ex L.) Thell. & *Salsola tragus* L. *Erodium cicutarium* is the most abundant, dominant across tens of thousands of acres within and adjacent to *Muilla lordsburgana*’s habitat. *Schismus barbatus* is often codominant with *Erodium cicutarium* along the lower-elevation margin of this habitat, and in many other areas of northern Hidalgo County. *Eragrostis lehmanniana* is on the upper-elevation edge of *Muilla lordsburgana*’s domain, and is not dominant across large swaths of the landscape in northern Hidalgo County, as it is in southeastern Arizona. *Salsola tragus* is ubiquitous, and sometimes dominant in finer-textured soils or following herbicide treatments.

Two other invasive species, *Sisymbrium irio* L. and *Descurainia sophia* (L.) Webb ex Prantl, are established in similar habitat near, but perhaps not yet within, areas occupied by *Muilla lordsburgana. Sisymbrium irio* is in dense, but relatively small, patches in northern Hidalgo County. It is dominant across large areas of similar habitat in northeastern Cochise and southeastern Graham counties of adjacent Arizona. *Descurainia sophia* is present in only a few small patches, although some of these are dense.

Loss of perennial grasslands probably plays a key role in the spread and abundance of these invasive plants. In other areas, including those more typical of Sandy and Deep Sand ecological sites, removal of grasses allows shrubs to become dominant. In this case, it appears to be invasive annuals that have taken advantage of the situation.

The effect of these changes in vegetation on *Muilla lordsburgana* are unknown, but are not likely to be positive. There is little direct human impact in the area other than livestock grazing, though. Climate change may pose a threat, if *Muilla lordsburgana* is near the limits of its heat or drought tolerance. Large solar facilities have been considered near Lordsburg, and could be a severe threat if built within the narrow range of the species.

### Additional populations

Shallowly incised bajadas with deep, coarse, sandy soils derived from intrusive igneous rocks are rare in southwestern New Mexico and adjacent Arizona, and relatively easy to identify using aerial imagery and geologic maps. The areas most similar to those occupied by *Muilla lordsburgana* are: to the southeast for ±32 km along the south side of the Big Burro Mountains; near Granite Pass on both sides of the Little Hatchet Mountains; on the north side of the Dos Cabezas Mountains; and on both sides of the northern Organ Mountains. Other patches can be found on both sides of the southern Pinaleño Mountains; sporadically on the bajadas of the Peloncillo Mountains (e.g., near Granite Gap); on the east side of the Pyramid Mountains; on the south side of Grandmother Mountain; and on the west side of the Florida Mountains. During March 2017, and more extensively during March and very early April 2020, I visited most of these areas to search for additional populations (Fig. 6), without success. Most of these sites did have some of the uncommon species associated with *Muilla lordsburgana*. If there are additional populations to be found, they are probably further west. Coarse sands derived from intrusive igneous rocks are more common to the west. Many of the associated species are more common to the west. *Muilla coronata*, presumably the closest relative to *Muilla lordsburgana*, is far to the west.

**Figure 6.**
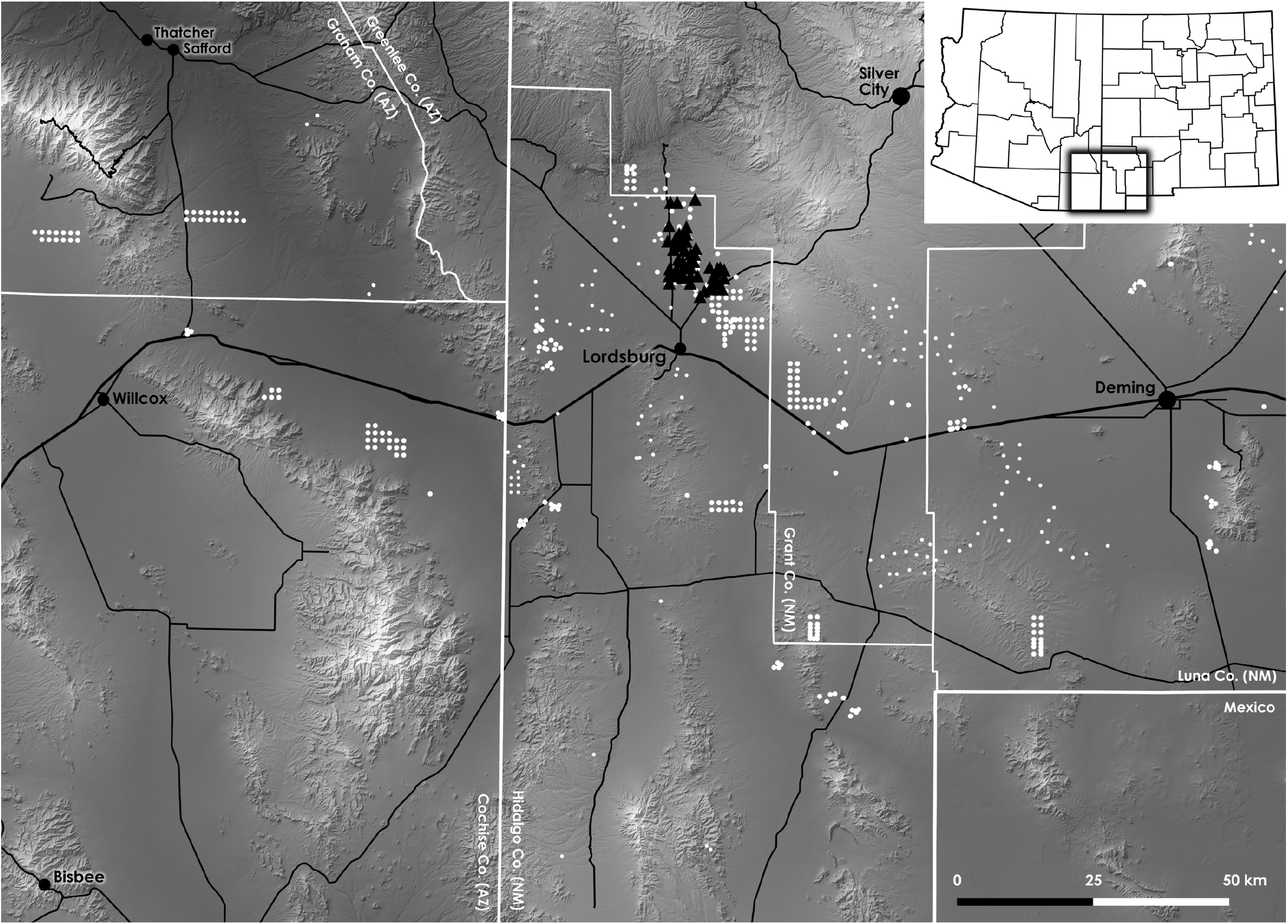
Distribution of *Muilla lordsburgana* (black triangles), areas visited while searching for *Muilla lordsburgana* (larger white circles), and other areas visited during the flowering period of *Muilla lordsburgana* (smaller white circles), when I was not looking for it but should have observed it were it present.

## Discussion

The discovery of *Muilla lordsburgana* highlights the unusual nature of soils derived from intrusive igneous rocks in southwestern New Mexico as well as the value of visiting underappreciated landscapes. In the course of field work on *Muilla lordsburgana* I encountered two species, *Calyptridium monandrum* (Nutt.) Hershkovitz and *Eriogonum thurberi* Torr., that have very rarely been recorded in New Mexico. Two more, *Logfia depressa* and *Stylocline sonorensis* Wiggins, are new records for the state. Specimen citations for these two are provided below. *Spermolepis organensis* G.L.Nesom, a narrow endemic described in 2012, occurs on very similar soils derived from quartz monzonite on the northeastern bajada of the Organ Mountains—including alongside the road to a popular hiking trail. These flat, open areas, bare much of the time and occasionally carpeted with *Erodium cicutarium &* its ilk after a wet winter, are not immediately appealing. Many botanists, myself included, have surely driven past all of these plants many times to arrive at some other destination.

The ecological dynamics of these habitats also deserve more attention. Areas where grassland has been lost without conversion to shrubland suggest greater complexity, and a less central role for shrubs, than the shrub encroachment paradigm (e.g., Grover & Musick 1990; Gibbens et al. 2005) would lead us to believe. Such areas turn out to be fairly extensive in southwestern New Mexico, once you have a reason to pay attention to them, and on a wider variety of soils than those discussed in this article.

### Logfia depressa

U.S.A. New Mexico. Grant County: Lordsburg Valley, between the Big Burro Mountains and Brockman Hills on the south side of Interstate 10, 4.8 miles north-northeast of Pigeon Hill, 32.21303°N 108.44719°W, 4480 ft., 4 Apr 2020, *Alexander 1639* (ARIZ, NMC, RENO, UNM). Hidalgo County: Granite Gap in the Peloncillo Mountains on the north side of NM Hwy. 80, 0.7 miles south of Preacher Mountain, 32.09408°N 108.96964°W, 4480 ft., 4 Apr 2020, *Alexander 1638* (ARIZ, NMC, RENO). Luna County: southwest of Grandmother Mountain, north-northwest of the Victorio Mountains, east of China Draw, and north of Interstate 10, 32.28892°N 108.16428°W, 4740 ft., 5 Apr 2020, *Alexander 1642* (NMC, RENO, UNM).

### Stylocline sonorensis

U.S.A. New Mexico. Grant County: Lordsburg Valley, between the Big Burro Mountains and Brockman Hills on the south side of Interstate 10, 4.8 miles north-northeast of Pigeon Hill, 32.21303°N 108.44719°W, 4480 ft., 4 Apr 2020, *Alexander 1640* (NMC, RENO). Hidalgo County: Granite Gap in the Peloncillo Mountains on the north side of NM Hwy. 80, 0.7 miles south of Preacher Mountain, 32.09408°N 108.96964°W, 4480 ft., 4 Apr 2020, *Alexander 1637* (ARIZ, NMC, UNM).

## Acknowledgements

I would like to thank Andrew C. Sanders, J. Chris Pires, David Griffin, and contributors to SEINet, CalPhotos, & iNaturalist for information regarding *Muilla coronata*, James D. Morefield for correcting my many misidentifications of minuscule members of Gnaphalieae, Guy L. Nesom for helpful comments on an earlier version of the manuscript, Matt Guilliams for assistance with *Pectocarya*, Roger A. Burks for tiny wasp identification, and J. Gregory Penn for assistance with associated plants & statistics.

## Data access

There are photographs related to this project in three categories: 1) Photographs of *Muilla lordsburgana* taken with a Canon 5D Mk. II, available as edited .jpg (at flickr.com/aspidoscelis and iNaturalist.org/observations/aspidoscelis) and RAW files; 2) photographs of habitat taken with a Canon 5D Mk. II, available as edited .jpg (at flickr.com/aspidoscelis) and RAW files; 3) photographs of *Muilla lordsburgana* taken with an iPhone SE (at iNaturalist.org/observations/aspidoscelis). All photographs, as well as coordinates and species lists for sites mapped in Figs. 5 & 6, are on Dryad: https://doi.org/10.5061/dryad.kkwh70s36.

This article presents the understanding of the author, who is not acting as a representative of the Bureau of Land Management.

The text of this copy of the article is, except for this sentence, identical to the version published as Journal of Semiarid Environments 1: 1-10 on 1 October 2021.

## Notes

### Competing Interest Statement

The authors have declared no competing interest.

## Literature cited

L.R. Abrams & R.S. Ferris, 1923. An illustrated flora of the Pacific States: Washington, Oregon, and California, vol. 1. Stanford University Press, Stanford, CA.

D.N. Cox, 1973. Soil survey of Hidalgo County, New Mexico. United States Department of Agriculture (U.S.D.A.), Soil Conservation Service, Washington, DC.

R.P. Gibbens, R.P. McNeely, K.M. Havstad, & R.F. Beck, 2005. Vegetation changes in the Jornada Basin from 1858 to 1998. Journal of Arid Environments 61(4): 651–668.

E. Gillerman, 1976. Mineral deposits of western Grant County, New Mexico. New Mexico Bureau of Mines and Mineral Resources, Bulletin 83.

E.L. Greene, 1888. New or noteworthy species II. Pittonia 1(4): 159–176.

H.D. Grover and H.B. Musick, 1990. Shrubland encroachment in southern New Mexico, U.S.A.: An analysis of desertification processes in the American southwest. Climate Change 17: 305–330.

J.W. Ingram, 1953. A monograph of the genera Bloomeria and Muilla (Liliaceae). Madroño 12: 19–27.

W.L. Jepson, 1909. A Flora of California, vol. 1. University of California Press, Berkeley, CA.

P.A. Munz, 1959. A California Flora. University of California Press, Berkeley, CA.

Natural Resources Conservation Service (U.S.D.A.), Jornada Experimental Range (U.S.D.A. Agricultural Resource Service), and New Mexico State University. 2020. Ecosystem Dynamics Interpretive Tool, https://edit.jornada.nmsu.edu/, accessed September 2020.

J.C. Pires & J.L. Reveal, 2002. Muilla. In: Flora of North America Editorial Committee, eds., Flora of North America North of Mexico, vol. 26, pp. 334–335.

J.C. Pires, 2017. Muilla. In: Jepson Flora Project (eds.) Jepson eFlora, http://ucjeps.berkeley.edu/eflora/eflora_display.php?tid=9541, accessed March & April, 2017.

QGIS Development Team. 2020. QGIS Geographic Information System 3.10 ‘A Coruña’. Open Source Geospatial Foundation Project. http://www.qgis.org/

J.R. Shevock, 1984. Redescription and distribution of Muilla coronata (Liliaceae). Aliso 10: 621–627

